# Prediction of Whole-Cell Transcriptional Response with Machine Learning

**DOI:** 10.1101/2021.04.30.442142

**Authors:** Mohammed Eslami, Amin Espah Borujeni, Hamid Doosthosseini, Matthew Vaughn, Hamed Eramian, Katie Clowers, D. Benjamin Gordon, Niall Gaffney, Mark Weston, Diveena Becker, Yuval Dorfan, John Fonner, Joshua Urrutia, Carolyn Corbet, George Zheng, Joe Stubbs, Alexander Cristofaro, Paul Maschhoff, Jedediah Singer, Christopher A Voigt, Enoch Yeung

## Abstract

Applications in synthetic and systems biology can benefit from measuring whole-cell response to biochemical perturbations. Execution of experiments to cover all possible combinations of perturbations is infeasible. In this paper, we present the host response model (HRM), a machine learning approach that takes the cell response to single perturbations as the input and predicts the whole cell transcriptional response to the combination of inducers. We find that the HRM is able to qualitatively predict the directionality of dysregulation to a combination of inducers with an accuracy of >90% using data from single inducers. We further find that the use of known prior, known cell regulatory networks doubles the predictive performance of the HRM (an R^2^ from 0.3 to 0.65). This tool will significantly reduce the number of high-throughput sequencing experiments that need to be run to characterize the transcriptional impact of the combination of perturbations on the host.

## Introduction

Cells enact complex dynamics in response to environmental and biochemical perturbations. The perturbation can have a widespread effect so as to alter the dynamics of the whole cell through cascading effects that span through a cell’s regulatory network. Combinations of the biochemical perturbations are thus not additive and can trigger complex responses such as heat shock ^1,2^, osmotic shock ^3,4^, or sudden shifts in nutrient availability ^5,6^. Given the complexity of these responses to these perturbations, prior studies in perturbed whole cell response examine the role of a specific well-known biophysical perturbation, for which there is a natural intuition or sensible alignment with known biophysical mechanisms ^7,8,9^. These experiments are carefully performed, driven by biophysical knowledge and hypothesis-based modeling.

A natural extension of this research is to examine how the whole cell responds to inputs that are foreign to the natural workings of the cell. Further, suppose that a cell was presented with multiple inputs, each of which lent biophysical insight, but the goal was to predict how the cell responded combinatorially to these inputs. The scale of experiments required to measure response in such a combinatorially large condition space is infeasible in terms of cost, labor, and time. In such a setting, there is great value in developing discovery-based approaches for spotlighting biophysical mechanisms using data-driven algorithms^10^, such as nonlinear modeling^11^ or machine learning^12^.

A domain that has grappled with modeling of combinatorially large condition spaces for prediction of specific cell response is that of drug combination/synergy prediction ^12,13^. Machine and deep learning techniques are widely used to model pharmacodynamic and pharmacokinetic parameters of a drug and identify biomarkers of drug response given a large corpus of drug and response features ^14–17^. The techniques used in these efforts require large training datasets that consist of specific cell responses to tens of thousands of drugs, a condition space that is often too large for high-throughput omics measurements, such as RNASeq, which provide insight into the whole-cell response. The ubiquity of high-throughput sequencing offers an opportunity to revolutionize the modeling of whole cell transcriptional response.

The ubiquity of high-throughput sequencing offers an opportunity to revolutionize the modeling of whole cell transcriptional response. A measure most often used to qualitatively and quantitatively assess a transcript’s response is its dysregulation as compared to a control. Differential expression analysis (DEA) is a standard bioinformatics technique that measures response to perturbations as compared to a control condition^18^. DEA conducts custom normalization, dispersion modeling, and Bayesian optimization to account for biological and experimental variability in the data. It quantifies the transcriptional response to a perturbation in terms of *fold-change* and measures its statistical significance. Data-driven prediction using machine learning of transcription to date, however, has been limited to expression level predictions from sequences or images ^19–20^. These techniques are prone to generalization errors that can arise from artifacts of normalization of counts data across experiments with combinatorically large condition spaces^21^.

In this paper we present the host response model (HRM), a machine learning model that can predict whole-cell transcriptional response to a combination of biochemical perturbations using transcriptional response data from single perturbations. Biochemical perturbations, in this context, amounts to inducing a cell with a chemical. The HRM combines high-throughput sequencing with machine learning to infer links between experimental context, prior knowledge of cell regulatory networks, and the RNASeq data to predict differential expression of a gene. The HRM was tested in two organisms, *Escherichia coli* MG1655 (*E. coli*) and *Bacillus subtilis* Marburg 168 (*B. subtilis*). *E. coli* is a well-studied and characterized Gram-negative bacteria that served as a proof of concept in the development of the HRM. *B. subtilis* is a well characterized and frequently used model organism for Gram-positive bacteria that was used to pressure test the HRM. For conciseness, the figures for *E. coli* are provided in Supplementary information.

## Results

### Training and Validation of a Machine Learning Model

In this study, we train and test a model per organism with embeddings of prior known transcriptional networks of the host cells to train three machine learning models for combinatorial prediction of DEA from single inducers **(Figure 1A**). The best performing model was selected using a validation dataset from two double-inducer conditions at two time points. Experiments are then conducted with all remaining inducer combinations at two time points to test the best performing model (in total 18 experimental conditions, 136 samples) **(Figure 1B).**

**Figure 1:**
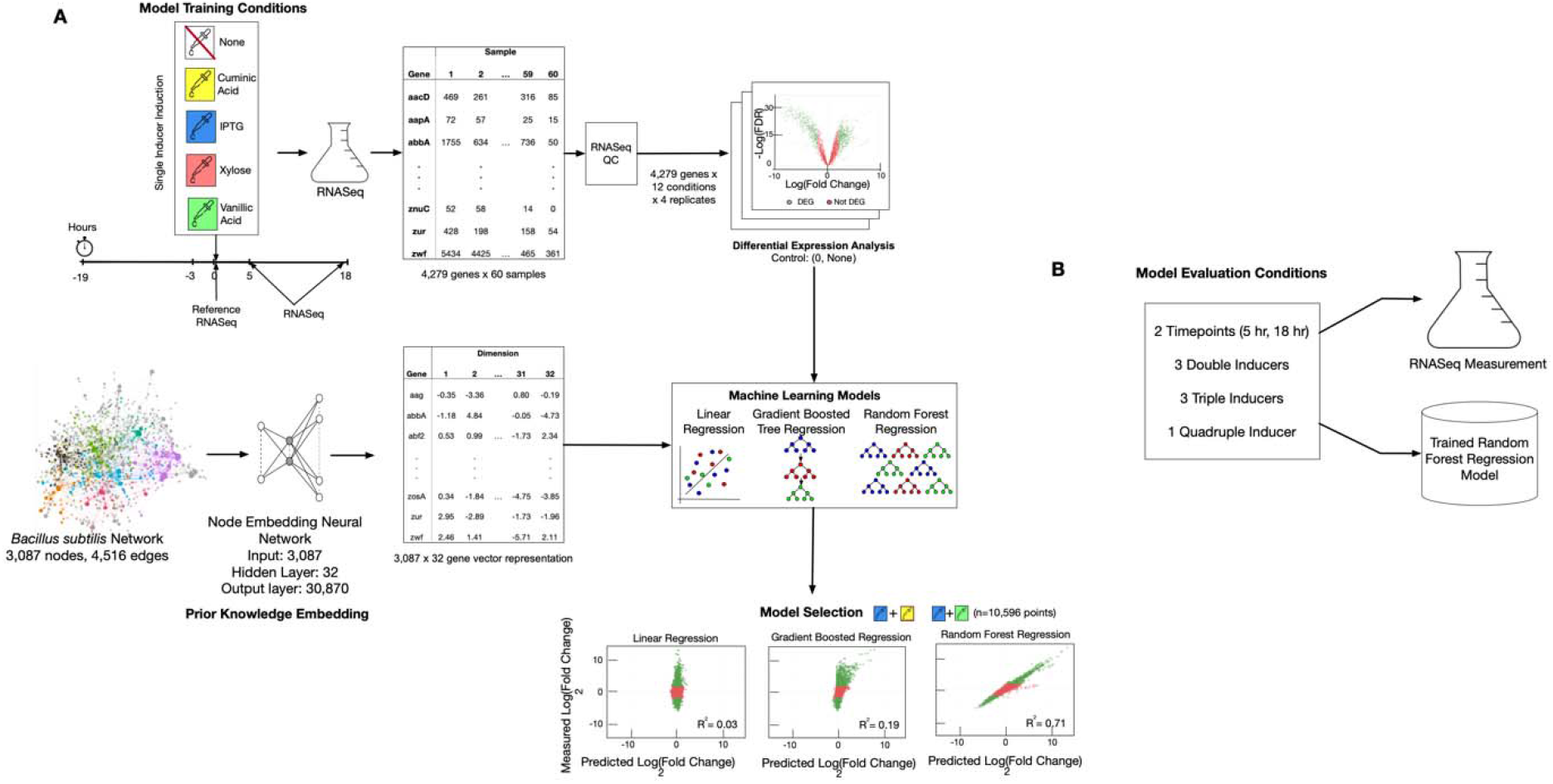
(A) Host response model experiment and knowledge integration for machine learning training and validation. For *B. subtilis*, RNASeq data was generated for single inducer conditions at each time point and passed through a configuration based differential expression analysis pipeline. Three machine learning models were validated with two held out pairs of inducer combinations. (B) The best model for the HRM is selected and tested with experiments run from all remaining combinations of inducers.

The HRM is formulated as a transcriptional dysregulation model trained with differential expression data and prior knowledge of gene networks of the host. The full set of conditions and samples collected for the training, validation, and test sets can be found in Supplementary Table 1 and a detailed description of the model can be found in the Methods section.

Training data for *E. coli* included used EColiNet^22^ as the prior gene network and experimental data that consisted of two inducers, Isopropyl β-d-1-thiogalactopyranoside (IPTG) and arabinose at four time points (5, 6.5, 8,18 hours). The test set was made up of the combination of the two inducers at all four time points. The non-induced, earliest time point (5 hours) was used as a control condition to measure host response using DEA for both training and testing data (Supplementary Figure 1). Training data for *B. subtilis* consisted of a transcriptional regulatory network^23^ with experiments at two phases of growth (log and stationary), and four inducers: IPTG, cuminic acid (CA), vanillic acid (VA), and xylose. The data was passed through a QC and DEA pipeline which removed low-quality samples and genes (Supplementary Table 2).

A challenge faced by the HRM is that the number of differential expression comparisons can have many factors, and thus, many design formulas. A Python-based configurable toolkit, which we call *omics_tools*, that parallelizes the execution of DEA for the large condition space was developed to address this challenge. The tool aggregates the outputs from the parallelized runs and combines all the data into a single unified dataset where each row represents a gene, it’s differential expression, statistical significance, and the condition for downstream machine learning. *Omics_tools* uses edgeR^24^ to conduct DEA across the set of design variables. A control condition of non-induced at the earliest time point was used to quantify the impact of induction and time. The same control condition was measured for all runs of the experiment that made up the training, validation and test set of data. The training corpus was formatted as follows: rows represented genes in each experimental condition, while columns consisted of features of the condition space, the node embedding features for the gene, and finally, the log fold change and associated statistical significance of the gene in the condition as compared to the control. The data can be found with the tutorial.

We find, surprisingly, a subspace representation of the individual responses enables prediction of response to combination of inputs. Qualitative performance of the model was measured as the number of dysregulated genes whose direction (up/down) was predicted correctly. Quantitative performance was measured with an R^2^ metric comparing predicted versus actual fold-changes on a logarithmic scale.

### HRM Predicts Transcriptional Response for E. coli

The first question to address was whether the set of differentially expressed genes had a large overlap between the train and test set. If so, then the task of machine learning would likely be trivial. We measured the Jaccard similarity between the set of genes of pairs of conditions to estimate the overlap (Supplementary Figure 2). The overlap between the conditions has a median of 0.2 with a standard deviation of 0.25, indicating a significant difference between the train and test differentially expressed genes (DEGs).

Three machine learning models were trained in two ways: using only genes present in the prior gene network versus using the whole transcriptome. To measure the impact of prior knowledge on the model, the best performing machine learning model with prior knowledge was selected and trained without prior knowledge to be used as a control. The criterion to label a gene as a DEG for *E. Coli* are genes with absolute log_2_(Fold Change) >1.1 and an FDR of <0.01. Qualitatively and quantitatively it was clear that machine learning could accomplish the task, but the impact of prior knowledge was marginal for *E. coli* (Table 1).

**Table 1:** *E. coli* results of qualitative and quantitative predictions for three different models using two training methods as compared to a control method for model selection.

| Model Name | Prior Networks<br>Used? | Training Method | Qualitative | Quantitative |
| --- | --- | --- | --- | --- |
| Gradient Boosted Regression | Yes | Genes only in network | 58.39% | 0.104 |
| Gradient Boosted Regression | Yes | Whole Transcriptome | 58.07% | 0.103 |
| Linear Regression | Yes | Genes only in network | 51.85% | 0.007 |
| Linear Regression | Yes | Whole Transcriptome | 51.73% | 0.006 |
| Random Forest Regression | Yes | Genes only in network | 90.20% | 0.887 |
| Random Forest Regression | Yes | Whole Transcriptome | 87.66% | 0.846 |
| Random Forest Regression | No | Control | 89.59% | 0.829 |

### Quantifying Prior Knowledge Impact for B. subtilis

Similar to the validation framework of *E. coli*, all single inducers and a subset of the double inducer data was used to train/validate the model (Table 2). Prior knowledge has a profound impact on the *B. subtilis* predictions showing that models trained and tested with genes only present in the prior network achieves >90% accuracy. Most interestingly, a model that does not use any prior knowledge of the host network achieved an R^2^=0.306, while one that used prior knowledge achieved R^2^=0.708, a 2.5x increase in performance.

**Table 2:**
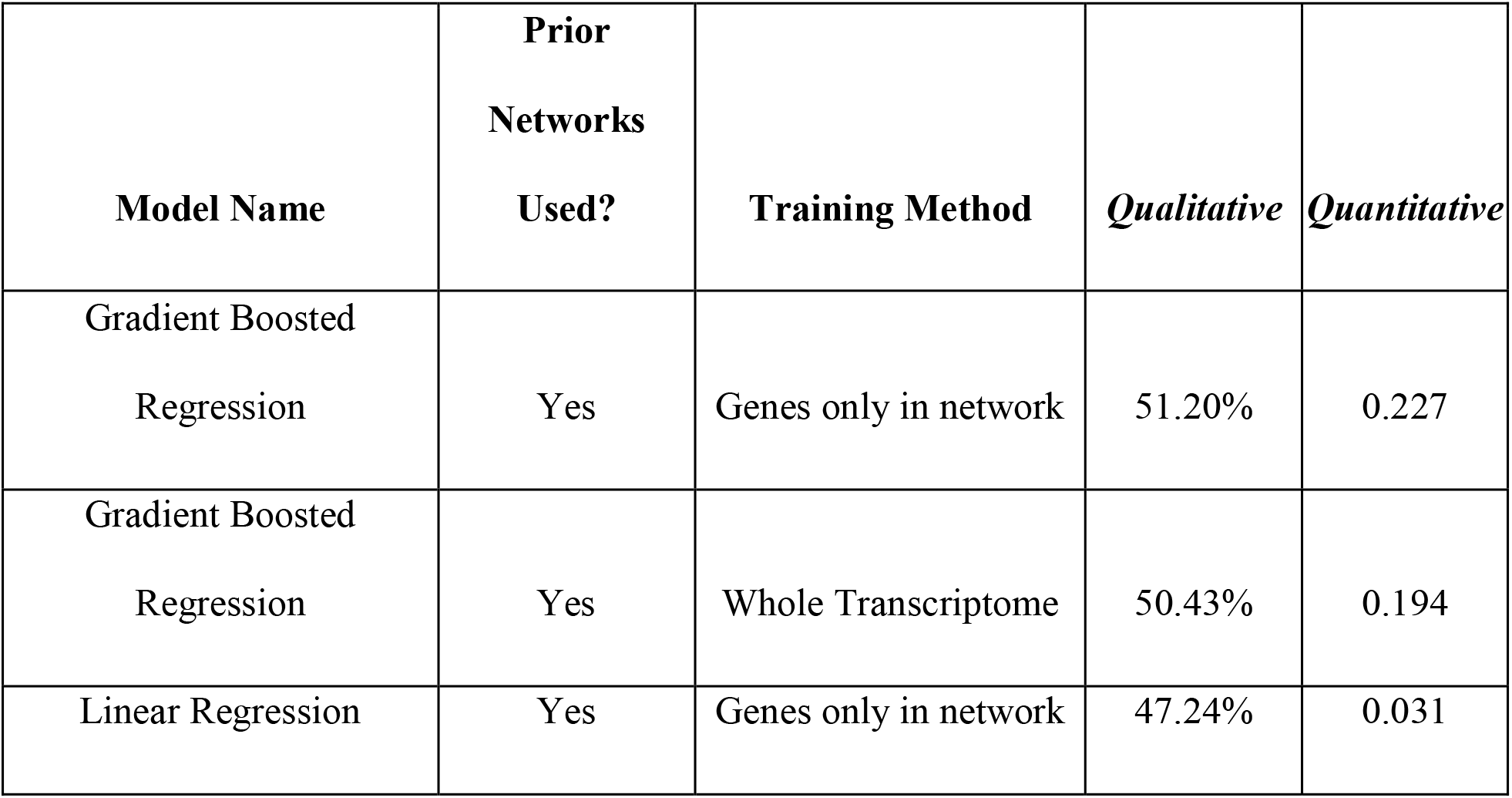
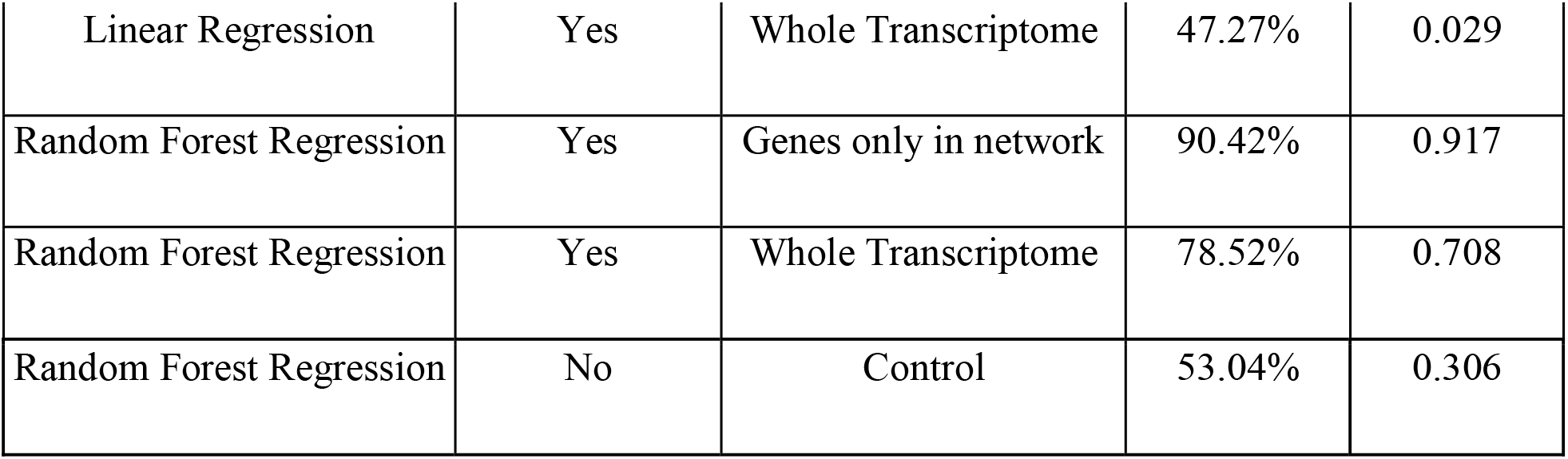
*B. subtilis* results of qualitative and quantitative predictions for three different models using two training methods as compared to a control method for model selection.

The discrepancy with *E. coli* can be explained by the heterogeneity of transcriptional response to the larger set of induction conditions. We computed a rank, or Spearman, correlation between the train and test induction conditions across all time points for both organisms. Specifically, this would be three comparisons for *E. coli* (double induced to none and single induced) and 39 comparisons for *B. subtilis* (single induced to all combinations). A distribution of the statistic is shown in Supplementary Figure 3. The larger distribution observed in the conditions of *B. subtilis* versus *E. coli* explains the impact of prior knowledge.

### Testing the HRM with All Inducer Combinations in B. subtilis

The best performing machine learning model, the random forest regressor, for the whole transcriptome was selected to test all remaining combinations. Specifically, predictions at remaining double, triple, and quadruple inducer conditions at the two time points were both qualitatively and quantitatively evaluated (Figure 2). Certain conditions could not be evaluated because there were less than two replicates that passed quality control (QC) (Supplementary Table 2 and Discussion for more details).

**Figure 2:**
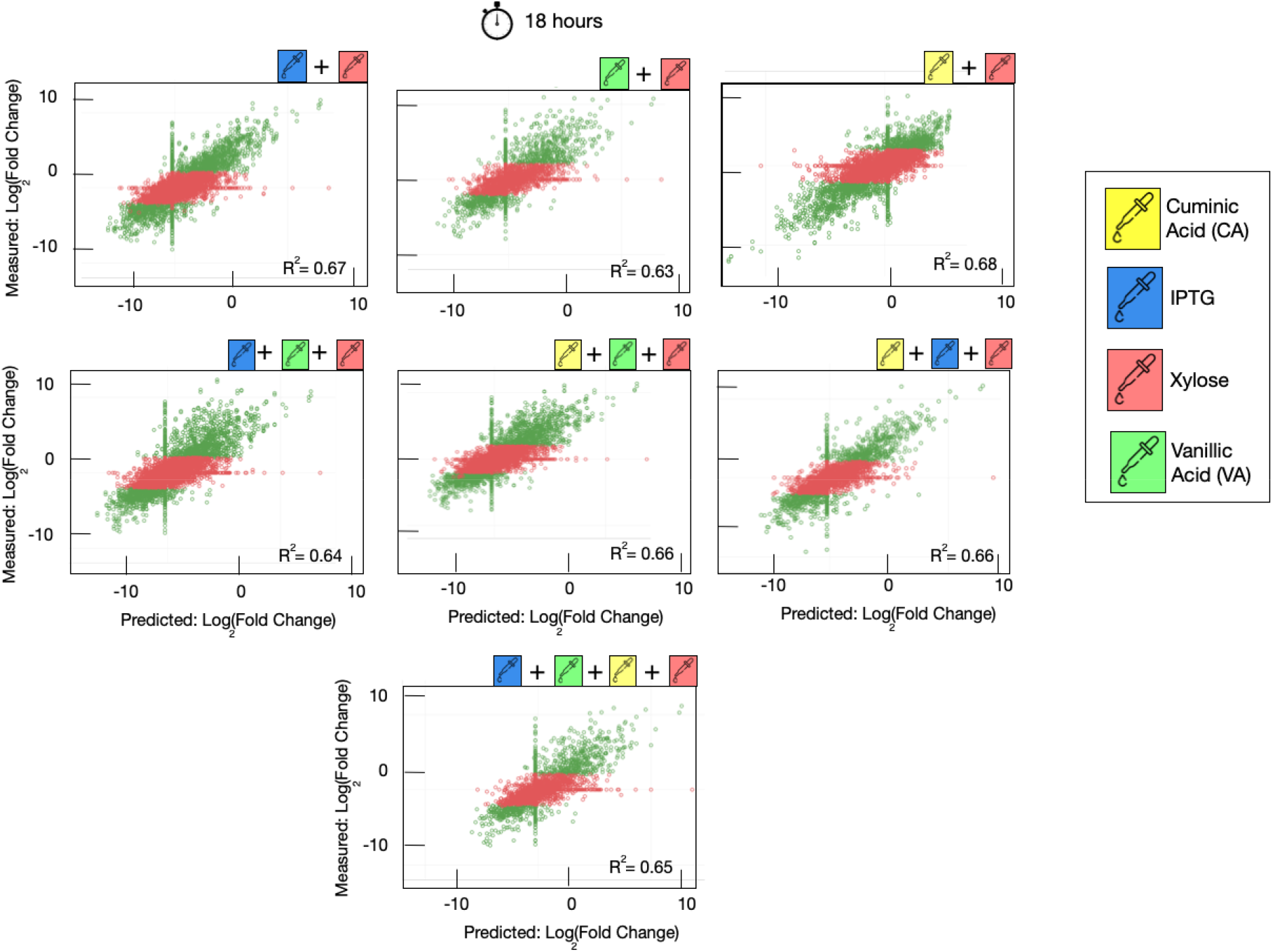
Predictions of transcriptional response -- log_2_(Fold change) -- for *B. subtilis* at 18 hours for over 2,000 genes. Conditions tested do not overlap with training and validation sets. Missing conditions are due to sample quality control. Red points are genes that are not differentially expressed while green genes are ones that are differentially expressed.

The model always maintained its performance within statistical error as the number of inducer combinations increased **(Figure 3A)**. Stationary phase predictions showed less variability and achieved >90% accuracy. The cells in log phase were of lower quality exhibiting lower OD and RNA integrity (Supplementary Figure 4) across all inducer conditions.

**Figure 3:**
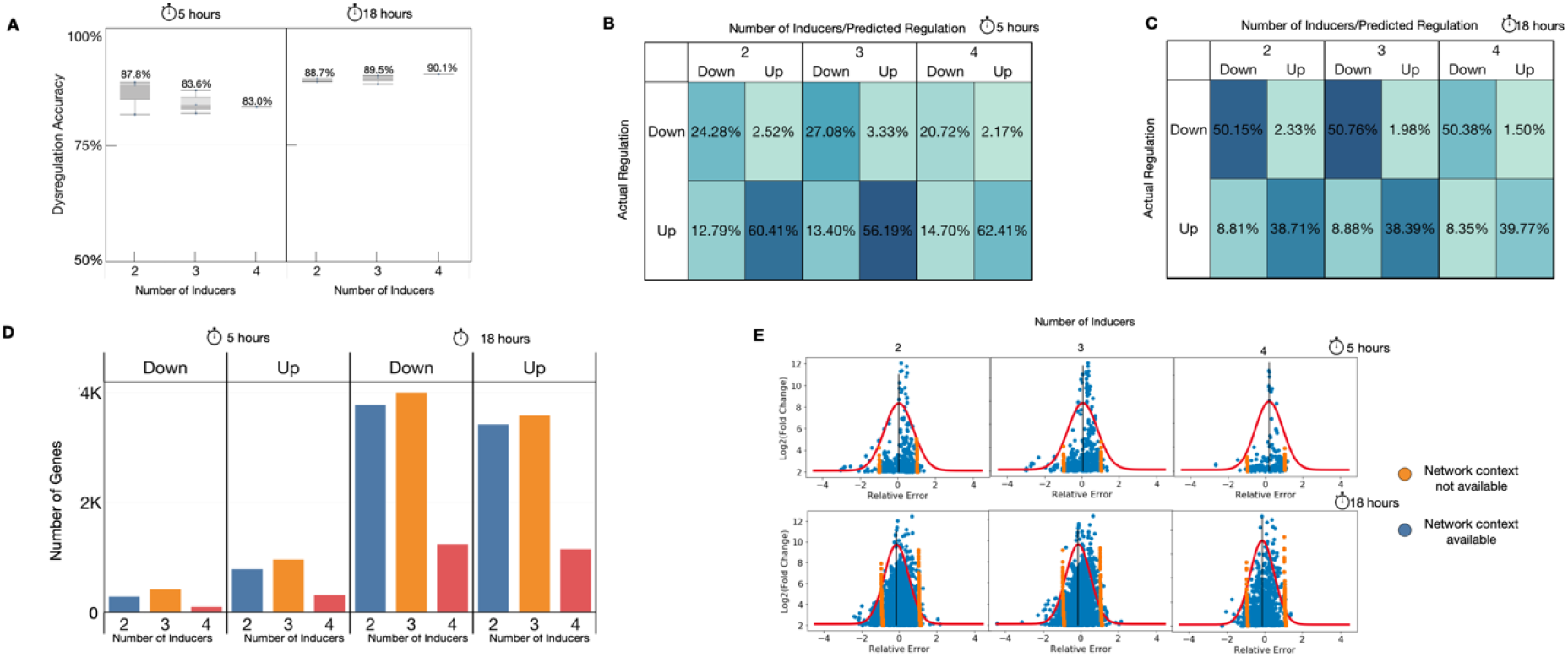
(A) Qualitative predictive performance of predictions on *B. subtilis* test set as it varies by the number of inducers. The performance stays within statistical error as the number of inducers increase. (B-C) Confusion matrices at 5 and 18 hours post-induction indicate that the model predicts more down-regulated genes as up-regulated. (D) The number of up and down regulated genes across inducers and time-points show 2x more up-regulated genes than down-regulated genes at 5 hours which indicates a large class imbalance. The distribution is more evenly balanced at 18 hours. (E) Quantitative error analysis shows most of the errors occur at |log_2_(Fold change)|<2.1 or when the gene is not present in the network. As a matter of fact, the gene’s not present in the network place an upper bound on the performance of the model.

Confusion matrices to assess the qualitative predictions show that the model had a more difficult time predicting up-regulated genes at both time points **(Figure 3B, 3C)**. Namely, there are more up-regulated genes predicted as down-regulated than there are down-regulated predicted as up-regulated. We should note that the log phase of growth had double the number of up-regulated genes as down-regulated ones (**Figure 3D**), commonly known as a class imbalance. This provides an opportunity to get more up-regulated genes incorrect at that phase. When the class is balanced, as in the stationary phase, the model’s predictions improve but hit an upper bound which will be explained in the quantitative assessment.

Quantitatively, the relative error is normally distributed with larger error at smaller dysregulations as those changes are harder to detect and predict. Genes with no network context available were mapped to a single point in the embedding space and so the model always predicted a constant value for all those genes **(Figure 3E)**. The network is composed of 2,608 genes from the total 4,266 genes in our reference strain. While this is >50% of the genome, genes with no network information make up only 25% of the DEGs. Even so, these genes make up the majority of qualitatively incorrect predictions and have greater than a single fold error (Table 3). This results in an upper bound on the performance of a model as the majority of the model’s errors are due to a lack of network context for certain genes.

**Table 3:** Number of incorrect predictions with greater than a single fold change error are primarily made up of genes with no embedding information.

| Number of Inducers in Test Conditions | Total number of predictions | Number of Incorrect Predictions due to absence of gene in network | Number of Incorrect Predictions due to presence of gene in network | Percent of Total Incorrect predictions due to absence of gene in network |
| --- | --- | --- | --- | --- |
| 2 | 3,765 | 1,016 | 134 | 88.4% |
| 3 | 4,050 | 1,092 | 142 | 88.5% |
| 4 | 1,240 | 313 | 41 | 88.4% |

## 4 Discussion

In this paper, we present a machine learning model enriched with features from prior, known transcriptional networks to qualitatively and quantitatively predict dysregulation of genes to a combination of induction conditions at two phases of growth. We showed that the use of a prior host network adds useful information to a model for it to make predictions of gene dysregulation for unseen combinations of induction conditions.

A natural next question one would have is how our predictions would impact analyses downstream of DEA, like enrichment analyses. These analyses help researchers gain mechanistic insight into gene lists generated from DEA ^30^. The goal here is to see how different the mechanistic insights would be if a researcher uses the HRM’s predictions from what they would have observed if they executed the experiments. To answer this question, we added annotations to the *B. subtilis* genes using SubtiWiki ^31^ to conduct enrichment analysis of predicted versus observed DEGs. No gene cluster file was publicly available to use standard enrichment tools. We created a gene cluster file using data from SubtiWiki (Supplementary Data 2) and used gene set enrichment analysis ^32,33^ to identify up and down regulated pathways for each condition (**Figure 4A)**. We evaluate the predictions by measuring the number of False Negatives (pathways that are down regulated, but we predict it is not down regulated), False Positives (pathways that are not down regulated, but we predict are down regulated), True Negatives (pathways that are not down regulated and we do not predict them to be down regulated), and, finally, True Positives (pathways that are downregulated and we predict them to be down regulated) (**Figure 4B)**. The same assessment is made for each test condition on the upregulated pathways (**Figure 4C).**As one would expect, the inducers do not regulate many pathways and the model correctly identifies most of those pathways (gray boxes). Every row is a level 2 category in SubtiWiki that is composed of multiple pathways. The number in each box represents the number of pathways in the category.

**Figure 4:**
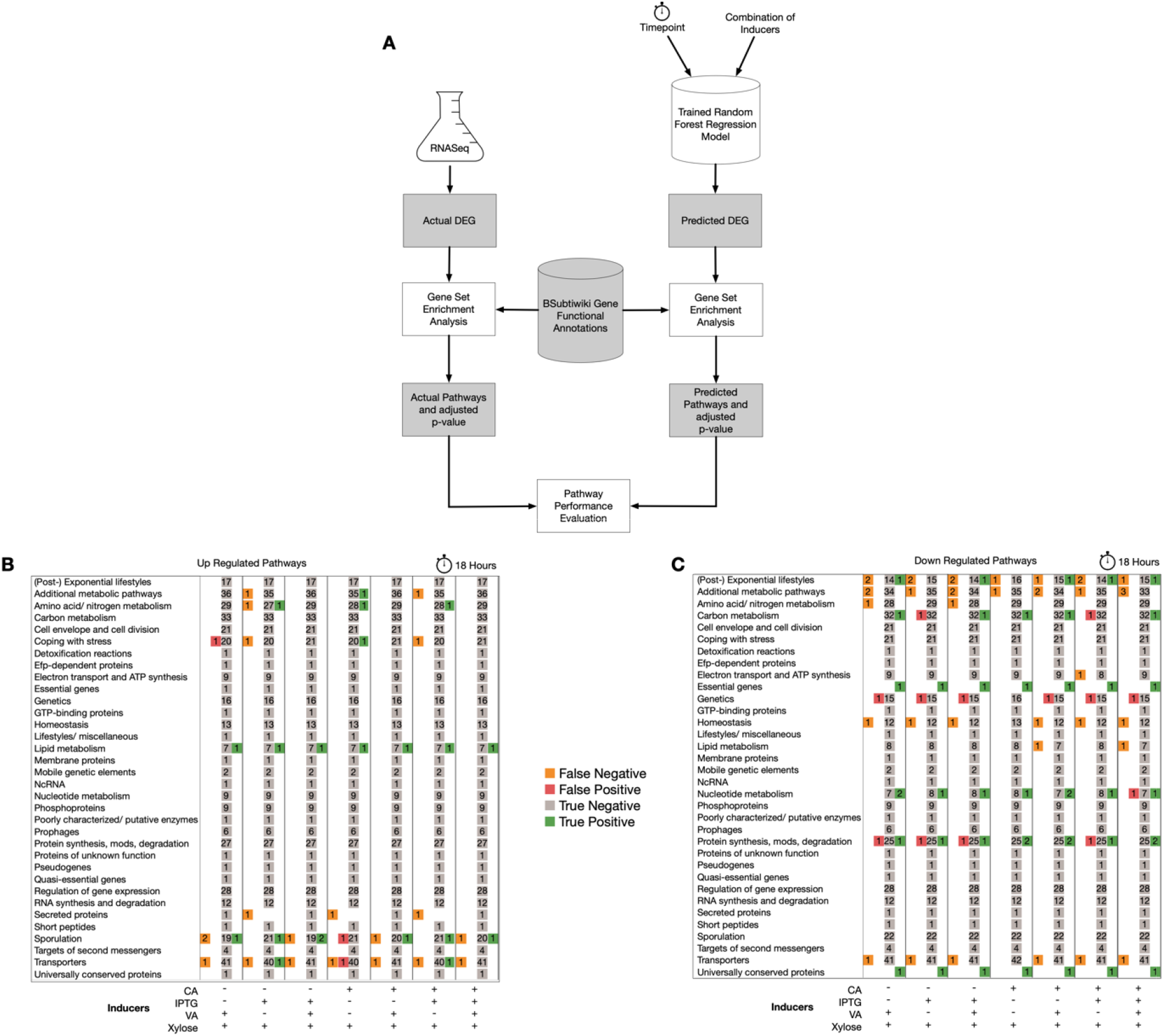
(A) Pathway analysis comparison of actual versus predicted set of DEGs using a gene cluster file derived from SubtiWiki. (B-C) Level 2 categories of annotations in the database composed of multiple down and up regulated pathways. The numbers inside each colored box indicates the number of pathways that were FP, FN, TP, TN. As expected, the inducers do not impact a majority of the pathways and the model accurately identifies those pathways.

Each pathway consists of a set of genes. Pathway analysis uses a statistical test (like a Fisher’s exact test) to identify dysregulated pathways by comparing the list of DEGs to each pathway's gene set. Thus, we should also check the precision of identifying the regulation of a pathway. This amounts to seeing how the predicted DEGs compare to the observed DEGs to identify if a pathway is up or down regulated. We picked a False Positive (FP), False Negative (FN), and True Positive (TP) sporulation pathway to assess the precision of the predictions (**Figure 5).** The FP was selected from the CA+xylose condition, while the FN and TP were selected from the IPTG+CA+VA+xylose condition, all at 18 hours. The genes that are in the bottom left quadrant of each plot are the ones that contribute to the pathway’s down-regulation status in both predicted and actual settings. The genes in the top left quadrant contribute to the down-regulation status of the pathway for predicted models but not in the observations. Finally, the genes in the bottom right quadrant contribute to the pathway’s down-regulation from the observations but not from the predicted model. The vertical set of orange genes are ones that are not present in the network and so the model predicts a constant for those expression values. The horizontal set of green genes are genes that did not pass QC and thus could not be validated. As indicated in the results section, these genes make up the majority of the genes that can be attributed to the enrichment errors. It is clear from this analysis that future work should consider jointly optimizing for differential expression as well as pathway inclusion.

**Figure 5:**
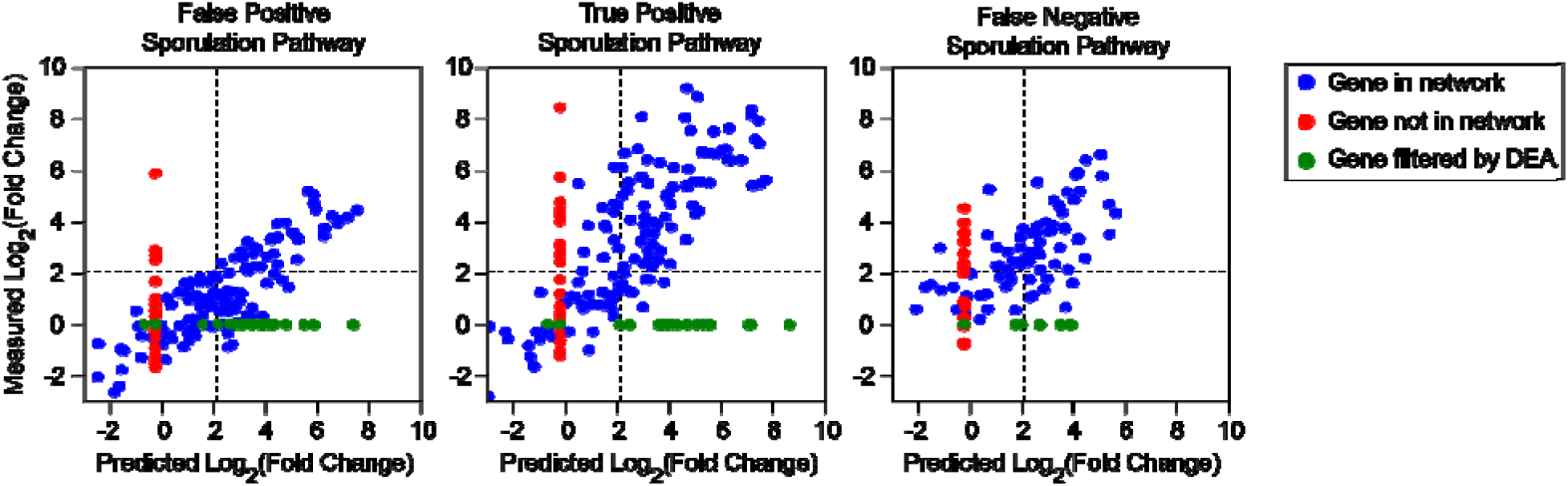
Predicted versus actual Log_2_(Fold change) of three pathways within the sporulation category that is a FP, TP, and FN. The genes in the top left quadrant contribute to the down-regulation status of the pathway for predicted models but not in the observations. Finally, the genes in the bottom right quadrant contribute to the pathway’s down-regulation from the observations but not from the predicted model. The set of orange genes are ones that are not present in the network and so the model predicts a constant for those expression values. The horizontal set of green genes are genes that did not pass QC and thus could not be validated.

## Methods

### Two Stage Learning Model to Predict Differential Expression

The goal to predict transcriptional response in a combinatorically large condition space from single conditions makes an end-to-end learning model with many free parameters underspecified and prone to generalizability errors^25^. To address this issue, we instead used a two stage learning process:

1. **Node embedding of Prior Knowledge**: We applied the node2vec algorithm to derive vector features from the network that could be used in the downstream learning task^26^. node2vec was selected because it is an unsupervised learning technique that balances depth and breadth first searches using a random walk to preserve both local and global connectivity structures of the genes in the network^27–29^. It can be parameterized by the length of the walks and the number of walks one takes from each node. A skip-gram model is then used to generate the vector embeddings in R^N^, in our case N was 32. We chose N=32 after an assessment of model predictions on the train/validation set sweeping N between 8, 16, 32, and 64. The performance was not statistically different and so we chose a parameter that was large enough to provide the model with degrees of freedom to generalize but small enough to ensure the model does not overfit. We should note that the network need not be for the exact strain being used, but should have significant genomic overlap. In our case, the overlap of EColiNet vs the genes in the MG1655 strains was 3818/4111 genes, and for *B. subtilis* was 2608/4266, which is >50% for each organism. Genes that were not present in the network were mapped to the origin in *R^32^*.
2. **Machine Learning Models:**We trained three machine learning models, a gradient boosted regressor, a linear regressor, and a random forest regressor, for their ability to predict the differential expression of a gene given the conditions of measurement and the derived network features. The models were trained on a regression task to minimize the error between predicted differential expression and the observed differential expression for a host’s response to single inducers. The induction conditions were one hot encoded to enable the representation of multiple induction conditions. Since there were so few timepoints for *E. coli* and *B. subtilis*, it was not treated as a continuous variable and was also one hot encoded.

The output predictions were evaluated using an R^2^ metric comparing predicted to actual differential expression in log_2_(Fold Change). We should note that only genes that were differentially expressed were used to measure R^2^ as those are the ones most significant in differential expression analysis. For *E. coli*, we defined a gene to be differentially expressed if it had an absolute magnitude of log_2_(Fold Change) >1.1 and FDR of <0.05, while for *B. subtilis* We defined a gene to be differentially expressed if it had an absolute magnitude of log_2_(Fold Change) >2.1 and FDR of <0.05.

### Sample Preparation and Processing

Wild type strains for B. subtilis (Bacillus Subtilis 168 Marburg) and E coli (E. coli K-12 MG1655) were cultured in M9 media consisting of 1X M9 media salts, 0.1mM CaCl2, 1X Trace Salts, 1mM MgSO4, 0.05mM FeCl3/0.1mM C6H8O7, 0.2% Casamino Acids, and 0.4% Glucose. The inducers used in this study were isopropyl β-D-1-thiogalactopyranoside (0.001 M), arabinose (25mM), vanillic acid (0.001 M), cuminic acid (0.0001 M), and xylose (1%).

Glycerol stocks were inoculated into M9 media in shake flasks, and the culture was grown overnight for 18h at 30◻°C and 1000rpm. The following day, cultures were diluted to OD 0.1 in fresh M9 media and grown in 96-well plates under the same conditions for 3 h. For induction, cells were diluted a second time to OD 0.05 in the presence of inducers. Plates were incubated at 30◻°C and 1000 rpm for 5 h and 18 h and cultured cells were harvested and fixed with either RNA protect (for E. coli) or methanol (B. subtilis).

Total RNA was extracted using Magjet RNA extraction kit (Thermo) according to manufacturer's instructions. RNA quality was assessed using Tapestation (Agilent). KAPA RNA Hyperprep kit (Roche) was used for ribosomal RNA depletion and Illumina compatible library preparation. Prepared library was loaded on a Illumina sequencer to generate 150bp paired end reads.

Raw RNA-seq data was trimmed and quality filtered with trimmomatic (v0.36), reads were aligned with bwa (v0.7.17). After alignment with bwa, the resulting sam files were sorted by PICARD tools (v2.18.15) function SortSam, and then AddOrReplaceGroups is run on the sorted sam. Gene-level quantification of counts was performed using the featureCounts function of Rsubread (v1.34.4).

### Samples and Transcript Quality Control for B. *subtilis*

A measure most often used to qualitatively and quantitatively assess a transcript’s response is its dysregulation as compared to a control. Differential expression analysis (DEA) is a standard bioinformatics technique that measures this response to perturbations as compared to a control condition ^19^. DEA conducts custom normalization, dispersion modeling, and Bayesian optimization to account for biological and experimental variability that is present in transcriptional counts data to quantify the transcriptional response to a perturbation and measure its statistical significance. While this method overcomes generalization errors that can arise from artifacts of normalization of counts data across experiments, it performs strict quality control (QC) at both the sample and gene level. These are listed below:

1. Sample QC = The significance tests to reject the null hypothesis of the differentially expressed genes require > 2 samples per condition. If this criterion is not met then DEA cannot be conducted.
2. Gene QC = DEA tools fits the Cox-Reid profile-adjusted dispersion to a set of normalized expressions across conditions. Genes that do not fit this profile are labeled as outliers and removed from DEA.

Three Boolean metrics were used to measure sample quality:

1. Number of mapped reads ≥ 500K
2. Count of all annotated genes ≥500K
3. Replicate correlation of a condition >0.9

If any of these metrics did not pass, the sample would be flagged as a low quality sample and not used for downstream analysis. For the *B. subtilis* experiments, we also collected OD600 measurements from a plate reader to correlate population with potential sample dropouts. We did not find a clear, discriminative correlation between this measurement and sample dropout for a condition, but the log phase measurements (timepoint 5.0) did have a lower OD on average and had twice as many samples that did not pass QC than stationary phase samples (timepoint 18.0). In conditions where only two replicates were available, differential expression analysis was not conducted and so those conditions could not be validated (Supplementary Table 2). All single inducer conditions for *B. subtilis* that passed QC were used to train the model. It should be noted, though, that if a sample passed QC, that did not mean all genes in that sample passed edgeR’s outlier detection method. edgeR fits normalized counts to a Cox-Ried dispersion model with a Bayseian optimization algorithm. A gene is removed by edgeR if it does not fit this dispersion model. While one can pass in a custom dispersion model per condition, we chose to use edgeR’s default, as development of custom noise models across the condition space was out of scope of this effort (Supplementary Table 3).

## Supporting information

B. subtilis gene matrix file for pathway analysis

## Funding

Any opinions, findings and conclusions or recommendations expressed in this material are those of the author(s) and do not necessarily reflect the views of the Defense Advanced Research Projects Agency (DARPA), the Department of Defense, or the United States Government. This material is based upon work supported by the Defense Advanced Research Projects Agency (DARPA) and the Air Force Research Laboratory under Contract No. FA8750-17-C-0231 (and related contracts by SD2 Publication Consortium Members).

## Data and Code Availability

The manuscript is accompanied with three code repositories that are fully documented with example python notebooks. The data for the publication is placed with the tutorials to ensure reproducibility of results.

1. A repository that includes the capability to train, validate, and test a machine learning model in a combinatorically large condition space. https://github.com/sd2e/CDM
2. A repository that includes a scaled, configurable differential expression analysis pipeline: https://github.com/SD2E/omics_tools.
3. A test-harness for machine learning models to make apples-to-apples comparisons of training and testing models: https://github.com/SD2E/test-harness

## Author Contributions

M.E., A. E. B, H. E., and E. Y. led the analysis plan, analysis, and design of experiments for E. coli and B. subtilis. H.D. also contributed to B. subtilis experiments. D. B., C. B., P. M, K. C., performed the experiments. J. U., M. W., M. V., G. Z., N. G., J. F., and J. S. designed and automated the data processing, quality control, and data/analysis collaboration infrastructure. M. E., G. Z, and A. C. developed, scaled, and automated the execution the differential expression analysis pipeline. M. E. and H. E. developed the machine learning test harness. M. E. and H. E. developed the Host Response Model Library. C. V., B.G., J.S., and Y.D are PIs that managed the program and provided technical guidance. M. E., A. E. B., and E. Y. prepared the manuscript.

## Competing Interests

The authors report no competing interests.

## Notes

https://github.com/sd2e/CDM

https://github.com/SD2E/omics_tools

https://github.com/SD2E/test-harness

### Competing Interest Statement

The authors have declared no competing interest.

