## Supplementary Text for "Prediction of Whole-Cell Transcriptional Response with Machine Learning"

### Supplementary Information

The host response model (HRM) is a machine learning model that is trained with differential expression analysis results from single inducers to predict response to combinations of inducers. Its predictive capabilities were evaluated with two organisms: Escherichia *coli* (E. *coli*) and Bacillus *subtilis* (B. *subtilis*). E. coli served as a proof of concept and a thorough test and evaluation was presented for B. *subtilis*. The conditions tested and their use in the model are listed below:

Supplementary Table 1: All experimental conditions used in this study.

| **Organism** | **Model Use** | **Inducers** | **Timepoints** | **Replicates** | **Total Samples** |
| --- | --- | --- | --- | --- | --- |
| E. *coli* | Training | None, IPTG, Arabinose | 5, 6.5, 8, 18 hours | 8 | 96 |
| E. *coli* | Testing | IPTG+Arabinose | 5, 6.5, 8, 18 hours | 8 | 32 |
| B. *subtilis* | Training | None, IPTG, Cuminic Acid (CA), Vanillic Acid (VA), Xylose | 0 (None only), 5, 18 hours | 4 | 44 |
| B. *subtilis* | Validation | IPTG + VA,  IPTG + CA | 5, 18 hours | 4 | 24 |
| B. *subtilis* | Test | None,  IPTG + Xyl,  CA + Xyl,  VA + Xyl,  IPTG + Xyl + CA,  IPTG+Xyl+VA,  VA+Xyl+CA,  IPTG+CA+VA,  IPTG + CA + VA +Xyl | 0 (None only), 5, 18 hours | 8 | 136 |

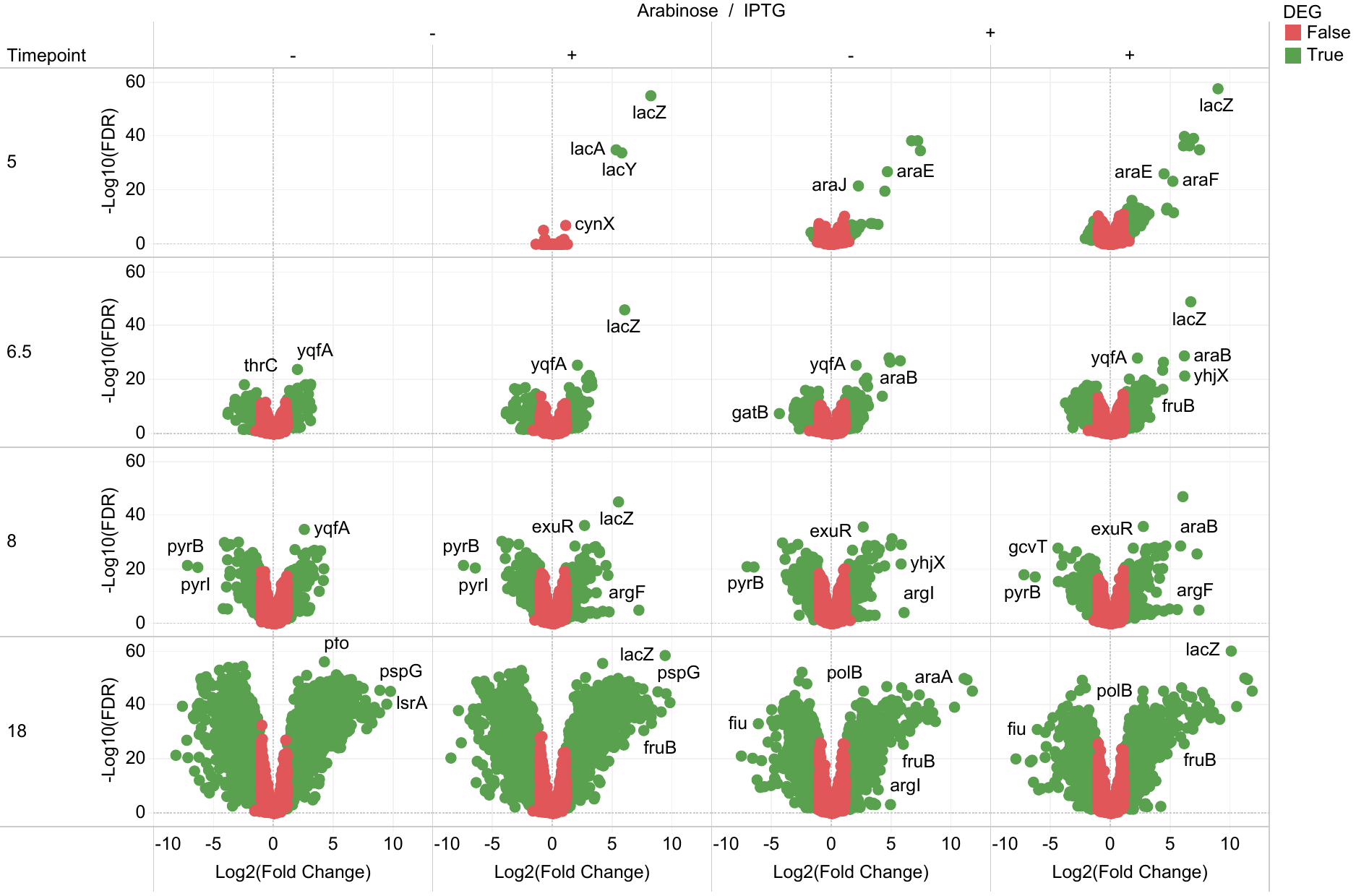

Supplementary Figure 1: Differential expression training set (left three columns) and test set (right column) for E. coli. +/- indicates presence or absence of inducer.

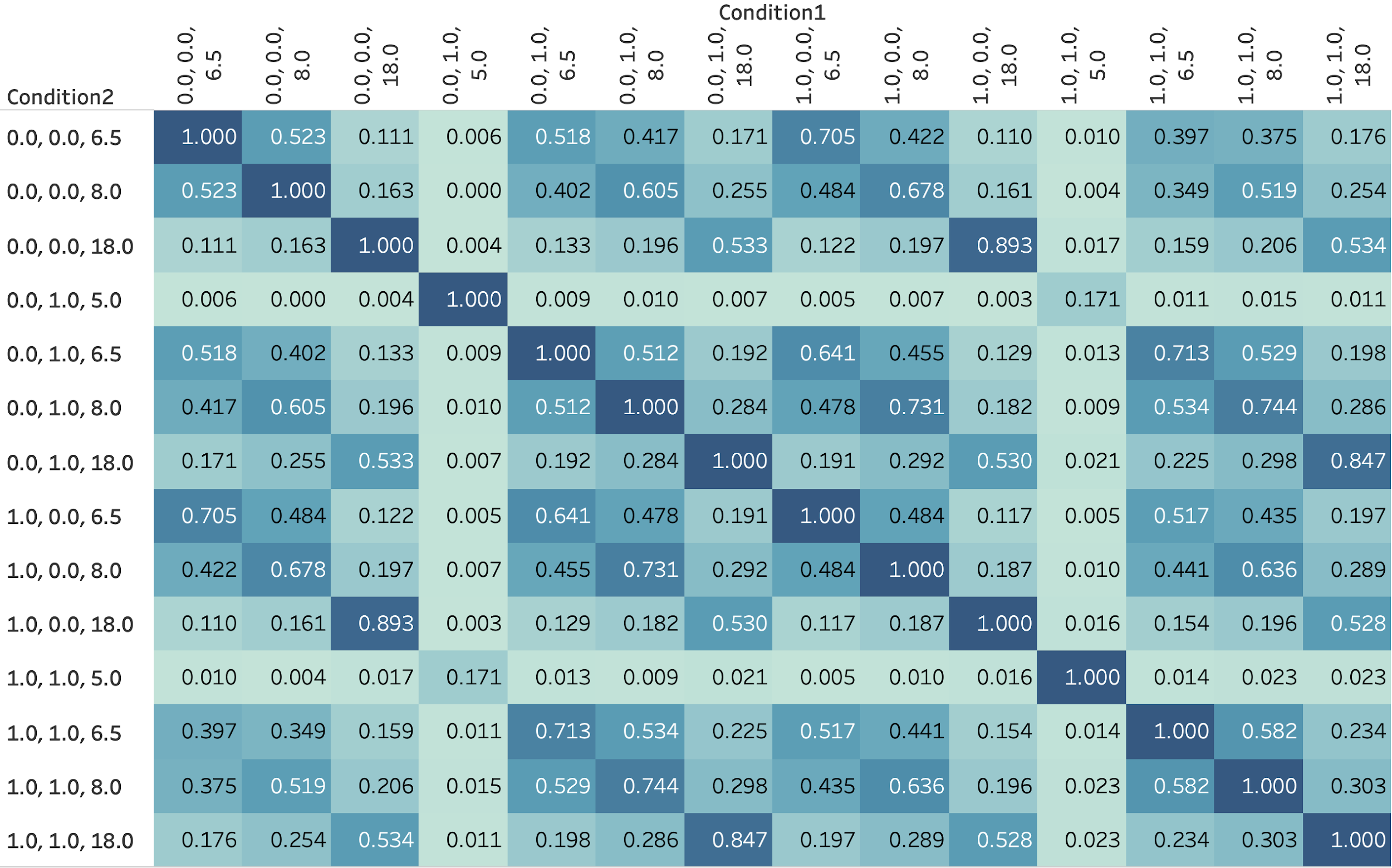

Supplementary Figure 2: Overlap of DEGs between the different conditions (1 is maximal overlap). Focus should be on the last four columns which capture the overlap between test conditions and the training conditions. The axes are tuples represented by (IPTG, arabinose, timepoint). For example, (0,0,6.5) refers to the condition where IPTG and arabinose are not present at 6.5hrs post induction.

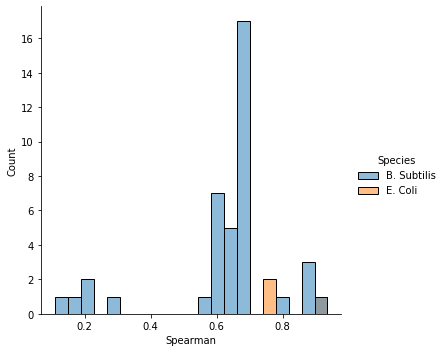

Supplementary Figure 3: Spearman correlation of log_2_(Fold change) distribution for E. *coli* and B. *subtilis*. The bar to the right is gray because there is an overlap of blue/orange in that bin.

Supplementary Table 2: Experimental test conditions with the number of replicates that failed (False) and passed (True) QC. A “0” indicates that the inducer was absent while a “1” indicates that the inducer was present.

| **Timepoint** | **Xylose** | **IPTG** | **CA** | **VA** | **Pass QC?** | **Num. Replicates** | **DEA Computed?** |
| --- | --- | --- | --- | --- | --- | --- | --- |
| Ctrl | 0 | 0 | 0 | 0 | TRUE | 9 | Yes |
| Ctrl | 0 | 0 | 0 | 0 | FALSE | 3 |  |
| 5 | 0 | 0 | 0 | 0 | TRUE | 4 | Yes |
| 5 | 0 | 0 | 1 | 1 | TRUE | 5 | Yes |
| 5 | 0 | 0 | 1 | 1 | FALSE | 3 |  |
| 5 | 0 | 1 | 1 | 1 | TRUE | 3 | Yes |
| 5 | 0 | 1 | 1 | 1 | FALSE | 5 |  |
| 5 | 1 | 0 | 0 | 1 | TRUE | 6 | Yes |
| 5 | 1 | 0 | 0 | 1 | FALSE | 2 |  |
| 5 | 1 | 0 | 1 | 0 | TRUE | 4 | Yes |
| 5 | 1 | 0 | 1 | 0 | FALSE | 4 |  |
| 5 | 1 | 0 | 1 | 1 | TRUE | 6 | Yes |
| 5 | 1 | 0 | 1 | 1 | FALSE | 2 |  |
| 5 | 1 | 1 | 0 | 0 | TRUE | 4 | Yes |
| 5 | 1 | 1 | 0 | 0 | FALSE | 4 |  |
| 5 | 1 | 1 | 0 | 1 | TRUE | 5 | Yes |
| 5 | 1 | 1 | 0 | 1 | FALSE | 3 |  |
| 5 | 1 | 1 | 1 | 0 | TRUE | 4 | Yes |
| 5 | 1 | 1 | 1 | 1 | TRUE | 6 | Yes |
| 5 | 1 | 1 | 1 | 1 | FALSE | 2 |  |
| 18 | 0 | 0 | 0 | 0 | FALSE | 4 | No |
| 18 | 0 | 0 | 1 | 1 | FALSE | 8 | No |
| 18 | 0 | 1 | 1 | 1 | FALSE | 8 | No |
| 18 | 1 | 0 | 0 | 1 | TRUE | 4 | Yes |
| 18 | 1 | 0 | 0 | 1 | FALSE | 4 |  |
| 18 | 1 | 0 | 1 | 0 | TRUE | 7 | Yes |
| 18 | 1 | 0 | 1 | 0 | FALSE | 1 |  |
| 18 | 1 | 0 | 1 | 1 | TRUE | 5 | Yes |
| 18 | 1 | 0 | 1 | 1 | FALSE | 3 |  |
| 18 | 1 | 1 | 0 | 0 | TRUE | 6 | Yes |
| 18 | 1 | 1 | 0 | 0 | FALSE | 2 |  |
| 18 | 1 | 1 | 0 | 1 | TRUE | 7 | Yes |
| 18 | 1 | 1 | 0 | 1 | FALSE | 1 |  |
| 18 | 1 | 1 | 1 | 0 | TRUE | 3 | Yes |
| 18 | 1 | 1 | 1 | 0 | FALSE | 1 |  |
| 18 | 1 | 1 | 1 | 1 | TRUE | 5 | Yes |
| 18 | 1 | 1 | 1 | 1 | FALSE | 3 |  |

The condition at 18 hours with no Xylose always had all replicates fail. We hypothesize that this is due to a lack of sufficient carbon source at 18 hours (there was 0.43% dextrose in the media which likely was not sufficient).

We collected optical density (OD) measurements to measure the impact the inducers would have on the cells. We did this as a means to see if there was an impact by the inducer on cell growth which did not allow us to extract RNA.

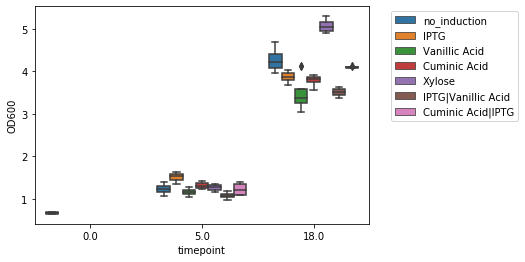

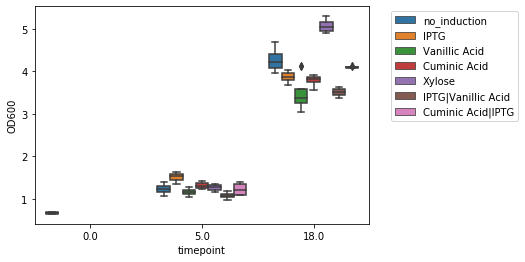

(a)

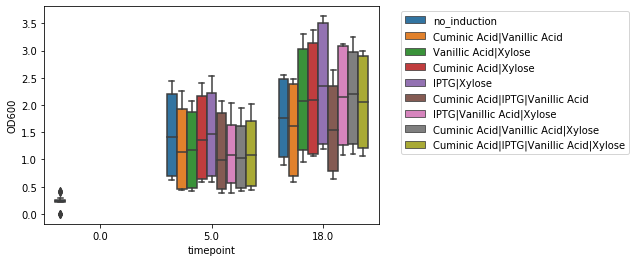

(b)

Supplementary Figure 4: OD600 measures for the (a) training/validation and (b) test conditions. OD<1 is often a flag for a potential RNASeq failure. Data for the training set was extremely well collected and so we expected high quality predictions from the model. Given the variability in the test set, however, means that many of those predictions may not have been validated.

Supplementary Table 3: Number of genes removed per induction condition across the two test runs.

| **Dataset** | **CA** | **VA** | **xylose** | **IPTG** | **Timepoint** | **Number of Genes Dropped out by edgeR** |
| --- | --- | --- | --- | --- | --- | --- |
| Test | 0 | 0 | 1 | 1 | 18 | 699 |
| Test | 0 | 1 | 1 | 0 | 18 | 695 |
| Test | 0 | 1 | 1 | 0 | 5 | 707 |
| Test | 0 | 1 | 1 | 1 | 18 | 695 |
| Test | 0 | 1 | 1 | 1 | 5 | 703 |
| Test | 1 | 0 | 1 | 0 | 18 | 905 |
| Test | 1 | 0 | 1 | 0 | 5 | 705 |
| Test | 1 | 1 | 1 | 0 | 18 | 688 |
| Test | 1 | 1 | 1 | 0 | 5 | 697 |
| Test | 1 | 1 | 1 | 1 | 18 | 666 |
| Test | 1 | 1 | 1 | 1 | 5 | 715 |
| Test Repeat | 0 | 0 | 0 | 0 | 5 | 624 |
| Test Repeat | 0 | 0 | 1 | 1 | 18 | 663 |
| Test Repeat | 0 | 0 | 1 | 1 | 5 | 653 |
| Test Repeat | 0 | 1 | 1 | 0 | 5 | 644 |
| Test Repeat | 0 | 1 | 1 | 1 | 18 | 572 |
| Test Repeat | 0 | 1 | 1 | 1 | 5 | 662 |
| Test Repeat | 1 | 0 | 1 | 0 | 18 | 599 |
| Test Repeat | 1 | 0 | 1 | 0 | 5 | 738 |
| Test Repeat | 1 | 0 | 1 | 1 | 18 | 590 |
| Test Repeat | 1 | 0 | 1 | 1 | 5 | 632 |
| Test Repeat | 1 | 1 | 0 | 0 | 5 | 803 |
| Test Repeat | 1 | 1 | 0 | 1 | 5 | 647 |
| Test Repeat | 1 | 1 | 1 | 0 | 18 | 570 |
| Test Repeat | 1 | 1 | 1 | 0 | 5 | 635 |
| Test Repeat | 1 | 1 | 1 | 1 | 5 | 675 |
